# Filopodial protrusion driven by density-dependent Ena-TOCA-1 interactions

**DOI:** 10.1101/2023.01.04.522504

**Authors:** Thomas C. A. Blake, Helen M. Fox, Vasja Urbančič, Adam Wolowczyk, Edward S. Allgeyer, Julia Mason, Jennifer L. Gallop

**Affiliations:** Wellcome/Cancer Research UK Gurdon Institute, University of Cambridge, Cambridge, UK; Department of Biochemistry, University of Cambridge, Cambridge, UK; Department of Genetics, University of Cambridge, Cambridge, UK

## Abstract

Filopodia are narrow actin-rich protrusions with important roles in neuronal development. The neuronally-enriched TOCA-1/CIP4 family of F-BAR and SH3 domain adaptor proteins have emerged as upstream regulators that link membrane interactions to actin binding proteins in lamellipodia and filopodia, including WAVE and N-WASP nucleation promoting factors and formins. Here, we demonstrate a direct interaction between TOCA-1 and Ena/VASP actin filament elongators that is mediated by clustered SH3 domain interactions. Using *Xenopus* retinal ganglion cell axonal growth cones, where Ena/VASP proteins have a native role in filopodia extension, we show that TOCA-1 localises to filopodia and lamellipodia, with a retrograde flow of puncta, and correlates with filopodial protrusion. Two-colour single molecule localization microscopy of TOCA-1 and Ena supports their nanoscale association. TOCA-1 clusters coalesce at advancing lamellipodia and filopodia and operate synergistically with Ena to promote filopodial protrusion dependent on a functional SH3 domain. In analogous yet distinct ways to lamellipodin and IRSp53, we propose that transient TOCA-1 clusters recruit and promote Ena activity to orchestrate filopodial protrusion.

## Introduction

Axonal growth cone navigation underlies accurate neuronal connectivity in the brain, guided by chemical and mechanical cues that are transduced to the cytoskeletal machinery to enable movement and turning (O’Connor *et al*., 1990; Koser *et al*., 2016). Cells move by combining protrusive and contractile forces with multiple morphological manifestations: lamellipodial driven motility, with sheet-like branched networks of actin (Rottner and Schaks, 2019), amoeboid motility where blebs of the cortex produced by hydrostatic pressure push the way through 3D enviroments (Bergert *et al*., 2015) and filopodial-driven motility.

Filopodia protrude through the dynamic growth and shrinkage of long, unbranched bundles of actin filaments, controlled by actin regulators at the filopodia tip (Mallavarapu and Mitchison, 1999; Applewhite *et al*., 2007). The actin bundle is surrounded by high negative curvature plasma membrane, stabilised by proteins that link curved plasma membrane to the cytoskeleton (Tsai *et al*., 2018). Control of filopodia protrusion, retraction and adhesion allows cells to transduce signals and transmit force to the environment via an adhesion-based molecular clutch (Chan and Odde, 2008; He *et al*., 2017). Filopodia promote tissue morphogenesis (Dent *et al*., 2007; Bischoff *et al*., 2021) and are a particular hallmark of axonal growth cones, where stochastic protrusion of dynamic filopodia has roles in transient adhesion to surfaces and promoting accurate movement (O’Connor *et al*., 1990; Dwivedy *et al*., 2007). How the processes of filopodia protrusion and remodelling between different actin-based architectures are controlled is important for understanding the mechanistic and genetic basis of disorders including cancer metastasis, intellectual disabilities and autism (Truesdell *et al*., 2015; Hu *et al*., 2016; Gouder *et al*., 2019; Wit and Hiesinger, 2022).

Within filopodia, the Ena/VASP family of proteins have been identified as major actin regulators underlying protrusion (Bear *et al*., 2000; Dwivedy *et al*., 2007; Barzik *et al*., 2014; Boyer *et al*., 2020; Damiano-Guercio *et al*., 2020). In lamellipodia, Ena/VASP activity determines the spacing between Arp2/3 complex branches and thus force generation on the membrane (Bear *et al*., 2002). At filopodia, Ena and VASP are localised to the growing tip, where their presence correlates with new actin monomers being incorporated into the elongating bundle (Applewhite *et al*., 2007; Urbančič *et al*., 2017). Biochemically, Ena and VASP are processive actin elongating proteins requiring G-actin binding activity, F-actin binding activity and oligomerisation for their function (Breitsprecher *et al*., 2008; Harker *et al*., 2019). They are subject to layers of regulation by a cyclical hierarchy of ubiquitination (Menon *et al*., 2015; Boyer *et al*., 2020) and by interactions with membrane adaptor proteins. These include IRSp53 and lamellipodin, sensors of negative membrane curvature and PI(3,4)P_2_ respectively, that direct the recruitment of Ena/VASP to clusters at the plasma membrane (Disanza *et al*., 2013; Montaño-Rendón *et al*., 2022), though neither are essential for Ena/VASP recruitment (Pokrant *et al*., 2022). Initiation pathways of filopodia have thus been proposed based on IRSp53 deformation of the membrane and recruitment of VASP (Tsai *et al*., 2022) and dynamic lamellipodin/VASP complexes that grow to a defined stoichiometry for controlled and productive protrusion (Cheng and Mullins, 2020).

The neuronally-enriched membrane adaptor protein TOCA-1 (also known as FNBP1L) and paralog CIP4 (present in mammals though not in *Xenopus*), are implicated in lamellipodial, filopodial and neurite formation in conjunction with N-WASP and WAVE yet their exact role has been elusive (Kakimoto *et al*., 2006; Bu *et al*., 2009; Fricke *et al*., 2009; Saengsawang *et al*., 2012, 2013). The actin polymerisation stimulating activity of TOCA-1 was first identified biochemically within *Xenopus* egg extracts (Ho *et al*., 2004) where N-WASP was discovered as a key binding partner, alongside a weak interaction with GTP-bound Cell division cycle 42 (Cdc42•GTP) (Ho *et al*., 2004; Watson *et al*., 2016). TOCA-1 also interacts with diaphanous-related formin 2 and orthologs (Aspenström *et al*., 2006) and CIP4 with WAVE in *Drosophila* (Fricke *et al*., 2009). To date, TOCA-1/CIP4 have been implicated in regulating actin polymerisation during endocytosis and endosomal trafficking (Fricke *et al*., 2009) but also in filopodia, with possible connections proposed between these processes (Bu *et al*., 2009; Nozumi *et al*., 2017; Gallop, 2020).

Within a cell-free model of filopodia-like structures that uses *Xenopus* egg extracts, TOCA-1 is one of the first proteins to be recruited to sites of actin polymerisation, and the intensities of TOCA-1 and Ena at the *in vitro* tip complexes are correlated with each other (Lee *et al*., 2010; Dobramysl *et al*., 2021). Ena and VASP are distinct in that Ena can be recruited to sites of TOCA-1 clustering prior to actin polymerisation, whereas VASP requires F-actin (Dobramysl *et al*., 2021). CIP4 and Ena (Mena, mouse Ena) have previously been seen to localize in mouse cortical neurons however it has not been determined how the interaction is controlled compared with other binding partners, and the dynamic implications of CIP4 or TOCA-1 enrichment on filopodial protrusion.

To study a cell type where profuse, long filopodia naturally form, we employed *ex vivo* dissected primary *Xenopus* retinal ganglion cells, which migrate from embryonic eye primordia along a laminin matrix to the tectum (Koser *et al*., 2016; Urbančič *et al*., 2017). A role for Ena (in *Xenopus* also referred to as Xena) has been characterised in filopodial protrusion and arborization in these cells (Dwivedy *et al*., 2007). We have mapped a direct biochemical interaction between TOCA-1 and Ena and identified a role for TOCA-1 in early filopodial extension and initiation. We show that the coincidence of TOCA-1 and Ena at lamellipodia and filopodia tips precedes forward filopodial movement in native filopodia and propose a density-dependent switch in TOCA-1 binding to Ena during the transition from lamellipodial to filopodial actin protrusion.

## Results

### TOCA-1 interacts with Ena directly *in vitro*, in a density dependent manner

Previously, we have examined the combinatorial correlations of 7 actin regulatory proteins with actin intensity and each other in a cell-free model of filopodia-like structures (Lee *et al*., 2010). TOCA-1 is one of the first proteins to arrive to the tip of filopodia-like structures and TOCA-1 levels correlate with those of other actin regulators including known interactors Diaph3 and N-WASP, as well as Ena and VASP (Lee *et al*., 2010; Dobramysl *et al*., 2021). We have measured the recruitment and correlations of Ena and VASP with filopodial dynamics in *Xenopus* retinal ganglion cell neurons, demonstrating the power of cross-correlation analysis to identify functional influence within redundant and stochastic molecular complexes at filopodia tips (Urbančič *et al*., 2017; Dobramysl *et al*., 2021).

While TOCA-1 interactions with N-WASP and Diaph3 have been reported previously, the association with Ena was unexpected, though conceivable through their SH3 and proline-rich regions (Fig. 1A). To determine whether TOCA-1 is capable of interaction with Ena and VASP, we covalently coupled recombinant SNAP-tagged *Xenopus tropicalis* TOCA-1, or SNAP alone, to benzylguanine derivatized magnetic beads and incubated them with *Xenopus* egg high speed supernatant extracts (Fox and Gallop, 2019). SNAP-TOCA-1 beads precipitated known interaction partners N-WASP, and Diaph3, plus Ena and VASP (Fig. 1B), confirming their status as possible interaction partners. To confirm that the SH3 domain was the site of interaction between TOCA-1 and Ena or VASP, each of the following constructs was expressed and coupled to magnetic beads: a point mutation in the SH3 domain which abolishes the interaction between TOCA-1 and N-WASP (W517K) (Ho *et al*., 2004), the F-BAR domain only, the F-BAR and HR1 domains, the F-BAR and HR1 domains with the subsequent linker region, or the SH3 domain alone (Fig. 1A). A functional SH3 domain is necessary and sufficient for Ena and VASP interactions (Fig. 1C).

**Fig 1.**
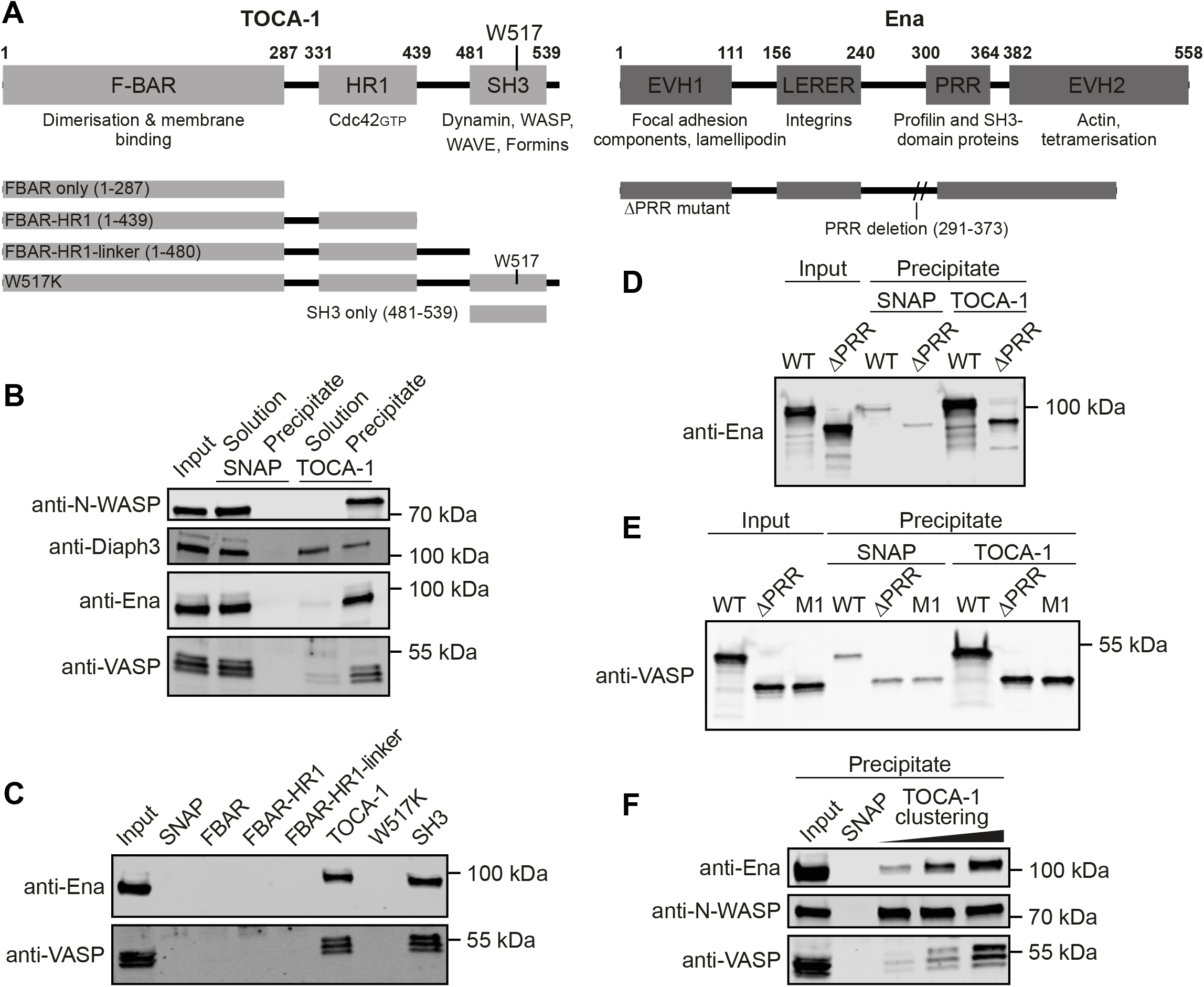
TOCA-1 interacts with Ena directly via SH3-PRR interactions in a density-dependent manner. (A) Schematic of TOCA-1 and Ena domain structures and mutations used in this study, numbered according to *Xenopus* sequences. (B) SNAP alone or SNAP-TOCA-1 coupled to benzylguanine magnetic beads and incubated with HSS, with Western blotting for known and candidate TOCA-1-interacting proteins. All the target proteins remained in solution when incubated with SNAP alone, while all were bound by SNAP-TOCA-1 with Ena and VASP interactions more marked than that of Diaph3. (C) The intact TOCA-1 SH3 domain is necessary and sufficient to bind Ena and VASP in HSS. (D) Purified TOCA-1 and Ena interact directly via the PRR of Ena. (E) Purified VASP and TOCA-1 interaction is not abolished by removal of the PRR or additional proline removal (M1 = ΔPRR + mutation of additional proline). (F) 1 nmol SNAP-TOCA-1 was coupled to 10, 50 or 100 μl SNAP-Capture beads followed by addition of beads up to a volume of 100 μl and an excess of SNAP protein in a second coupling step. HSS was added as in (B). Ena and VASP are bound more efficiently at higher densities of TOCA-1 while N-WASP binding efficiency is unaffected. Blots are representative of three repeats.

To confirm the interactions with TOCA-1 are direct and via the proline-rich regions (PRR) of Ena and VASP, purified recombinant Xenopus wildtype and ΔPRR proteins (Fig. 1A) were incubated with immobilized, purified SNAP-TOCA-1. Purified wildtype Ena and VASP bound directly to SNAP-TOCA-1 (Fig. 1D,E). Deleting the PRR of Ena reduced binding (Fig. 1D), however neither PRR deletion nor mutation of an additional proline within VASP meaningfully reduced binding (Fig. 1E). As the W517K point mutation in TOCA-1 eliminated VASP binding, this is a surprising result. VASP does not precipitate with SNAP beads alone (Fig. 1E) suggesting either that there are other possible candidate prolines within VASP or that there is a different type of molecular interaction sequence within VASP responsible for its interaction with TOCA-1. We note however that in our previous work, VASP behaved differently to Ena, with VASP being more dependent on polymerised actin for recruitment to *in vitro* filopodia-like structures and showing a lower level of correlation with TOCA-1 (Dobramysl *et al*., 2021).

The tetramerization of Ena and VASP is critical to their functions (Harker *et al*., 2019) whereas N-WASP is a monomer that undergoes clustering and dimerisation dependent on its interaction with SH3 domain-containing binding partners (Padrick and Rosen, 2010). One example is via TOCA-1, which has an F-BAR domain that forms both dimers and higher order oligomers on membrane surfaces (Frost *et al*., 2008). We therefore tested whether the interaction of TOCA-1 with N-WASP, Ena or VASP was sensitive to the presence of highly clustered TOCA-1, mimicked by immobilising SNAP-TOCA-1 and SNAP alone in different densities on the benzylguanine beads (Fig. 1F). While the same amount of N-WASP was bound by a fixed quantity of SNAP-TOCA-1 regardless of whether it was sparsely or densely coupled to beads, Ena and VASP showed a strong preference for a dense coupling of SNAP-TOCA-1 (Fig. 1F). Because the VASP interaction site with TOCA-1 could not be defined and it has reduced functional association with TOCA-1 within a multi-molecular environment compared to Ena (Dobramysl *et al*., 2021), we continued our study using Ena.

### TOCA-1 localises to RGC growth cone filopodia, dependent on a functional SH3 domain

TOCA-1 has previously been seen to localise to and stimulate filopodia in N1E-115 neuroblastoma cells when co-overexpressed with N-WASP (Bu *et al*., 2009). To determine whether TOCA-1 localises to filopodia in primary cells we microinjected RNA encoding mNeonGreen-TOCA-1 (mNG-TOCA-1) and GAP43-RFP membrane marker into the 4 cell stage of *Xenopus* embryos, or electroporated at stage 26-28 once eyes had formed, and microdissected eye explants at stage 32. After culturing *ex vivo* for 24 hours on laminin-coated glass dishes, RGC axons grow out from the eyes in a manner that is amenable to high speed, simultaneous two-colour total internal reflection fluorescence (TIRF) microscopy. RGC filopodia are used during navigation and arborization in development of the retinotectal system of Xenopus embryos (Dwivedy *et al*., 2007).

TOCA-1 formed distinct puncta with a striking localisation to filopodia tips, advancing lamellipodia and inwardly-moving puncta within the central domain (Fig. 2A; Video S1). A similar pattern of localisation was seen for human mNG-TOCA-1 transfected into human neuroblastoma SH-SY5Y cells (Fig. S1). Tracking individual TOCA-1 plasma membrane-localised puncta over four minute time-lapse videos revealed that 46% persisted at the plasma membrane, 34% moved into the central domain and 21% localised to filopodia, either at the tips or mobile within the shaft (data from 107 TOCA-1 puncta in 4 growth cones). Of newly-forming filopodia, 65% had TOCA-1 at the tip, while in 41% of filopodia TOCA-1 could be seen departing from the base of the filopodium during formation (data from 17 filopodia from 5 growth cones). In 24% of filopodia, no TOCA-1 was seen at any stage.

**Fig. 2.**
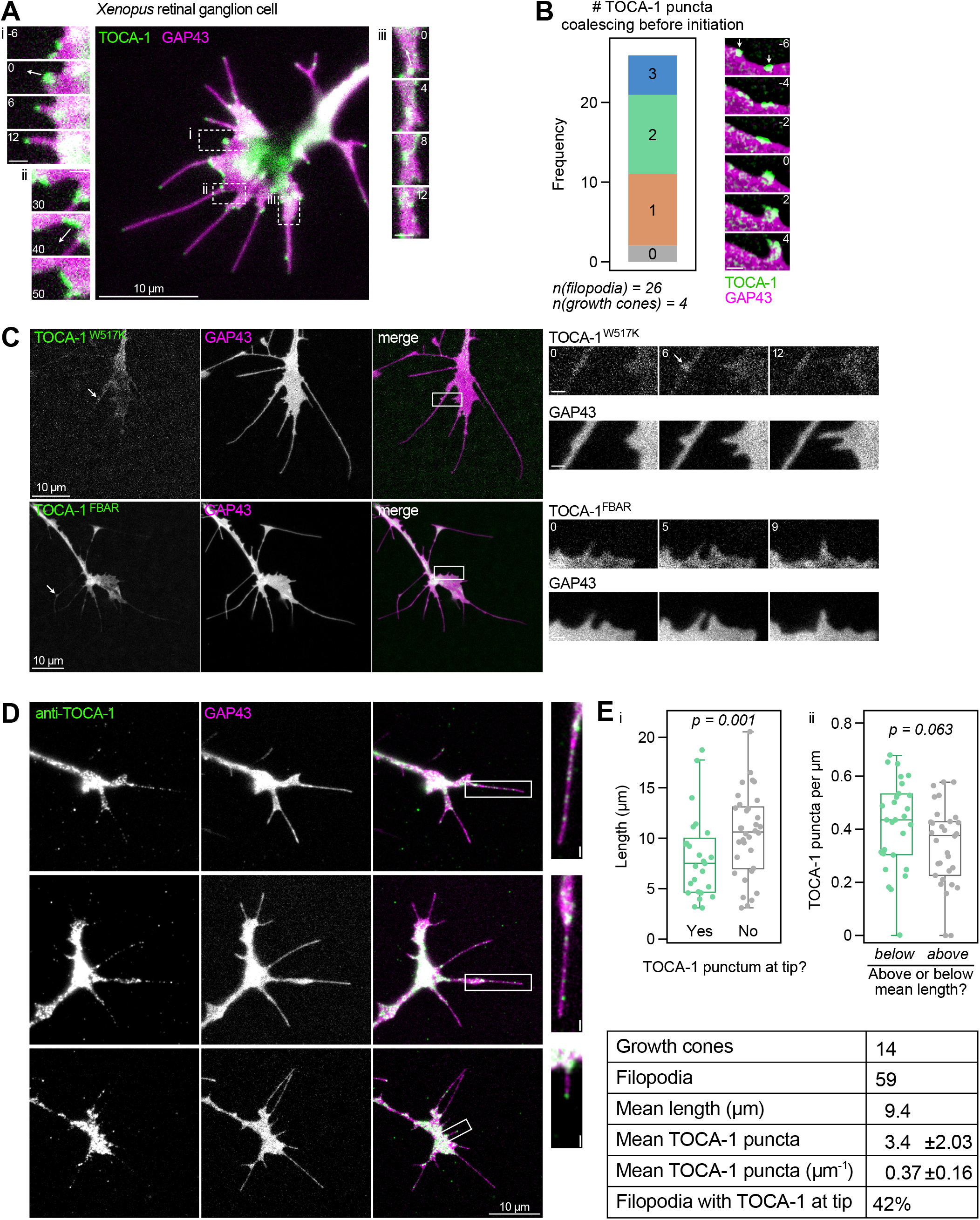
TOCA-1 localises to neuronal filopodia, dependent on a functional SH3 domain. (A) A *Xenopus* RGC axonal growth cone (Video S1) after micro-injection with mNG-TOCA-1 and the membrane binding region of GAP43-RFP, showing localisation of mNG-TOCA-1 to (i) filopodia, (ii) advancing lamellipodia and (iii) inwardly-moving puncta in the growth cone body. Arrows show direction of punctum movement. (B) Scoring of the number of distinct puncta of mNG-TOCA-1 observed to coalesce in the preceding few seconds at the site of future filopodium formation. Example montage (Video S2) shown with two distinct puncta indicated by arrows. Images de-noised with nd-safir. (C) Images showing RGC axonal growth cones expressing mNG-tagged TOCA-1 mutants (mNG-TOCA-1-W517K (*top*; Video S3) and TOCA-1-FBAR (*bottom*)), with arrows showing sites of moderate TOCA-1 enrichment, and (*inset*) areas of dynamic membrane showing no strong TOCA-1 enrichment. (D) Fixed RGCs expressing GAP43-RFP, immunostained with affinity purified anti-TOCA-1, showing on average 3.4 puncta of TOCA-1 per filopodium, in shafts and tips (see examples inset). (E) Filopodia (i) with a TOCA-1 punctum <1 μm from the tip were significantly shorter than those without (p = 0.001; Kruskal-Wallis test). (ii) Filopodia with below average length had a trend towards higher TOCA-1 puncta density (p = 0.063; Student’s t-test, two-tailed, equal variances). All scale bars are 1 μm unless indicated, time shown in seconds.

In 15/26 newly forming filopodia, 2-3 distinct puncta of TOCA-1 were observed to coalesce in the few seconds before filopodium formation, forming a single large punctum which often stayed at the tip, but in some cases split in two with fluorescence moving into the central domain while a punctum of TOCA-1 followed the protruding filopodium tip (Fig. 2B, Video S2). This is similar to previous observations for the lamellipodin-VASP complex, where splitting of filopodial initation complexes was implicated in maintaining lamellipodin-VASP stoichiometry (Cheng and Mullins, 2020).

To determine which domain of TOCA-1 is responsible for its localisation to filopodia, RGCs were prepared that express mNG-TOCA-1-SH3^W517K^ or mNG-TOCA-1-FBAR alone (Fig. 1A). Either removal or mutation of the SH3 domain results in diffuse, cytoplasmic localisation of TOCA-1, with no strong enrichment to filopodia, though weak enrichment was observed at swellings on the filopodia similar to the convoluted nodes associated with actin patches in dendritic filopodia (Fig. 2C, Video S3) (Galic *et al*., 2014). To confirm that endogenous TOCA-1 localises to filopodia we performed immunostaining with an affinity purified antibody raised in rabbit against full-length *Xenopus* TOCA-1 (Fig. S2). Similar to mNG-TOCA-1, immunostaining of endogenous TOCA-1 in RGCs shows localisation to the tips and shafts of filopodia (Fig. 2D shows 3 typical examples). Filopodia with TOCA-1 typically contained several puncta within the shaft, and the 42% with TOCA-1 present at the tips were on average shorter than those without, while those shorter than average tended to have a higher density of TOCA-1 puncta (Fig. 2E). Together, these results show that TOCA-1 is specifically enriched in active regions of lamellipodia and filopodia and it is dependent on a functional SH3 domain.

### TOCA-1 abundance correlates with filopodial initiation and tip movement

Because TOCA-1 localisation at tips seemed biased towards short filopodia, we performed timelapse TIRF imaging to determine its abundance at filopodia tips during the different phases of growth and quantified the tip movement and TOCA-1 intensities in filopodia expressing mNG-TOCA-1 (Fig. 3A,B). In some filopodia, it was evident that fluorescence transiently peaks at or shortly before extension (Fig. 3B).

**Fig. 3.**
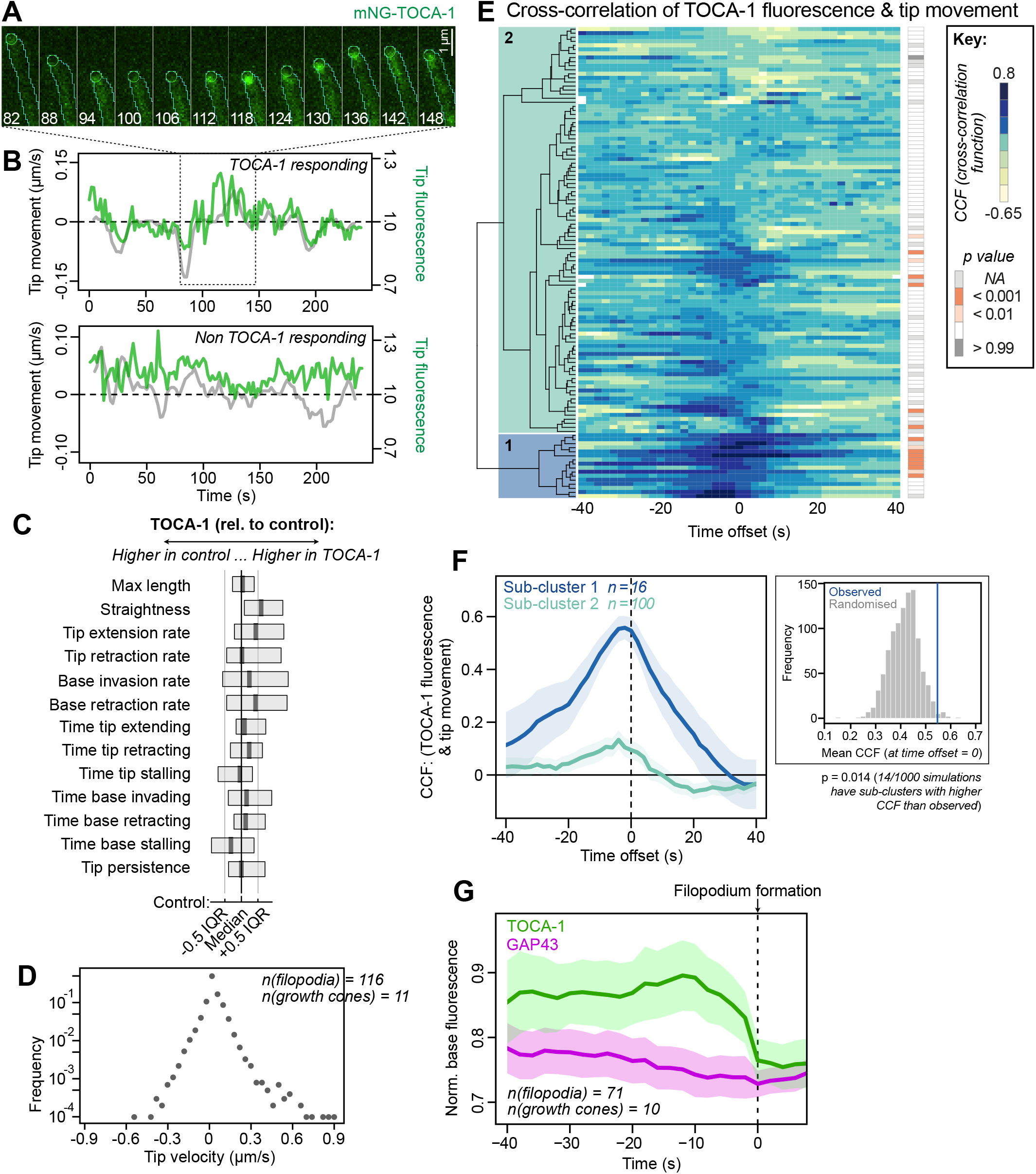
Fluorescence intensities and correlations of TOCA-1 with filopodial initiation and tip movement. (A) Montage of filopodia (cyan outline) and tip (green circle) segmented by Filopodyan, showing an increase in mNG-TOCA-1 fluorescence at the moment of forward movement. (B) Quantification of fluorescence intensity of two independent filopodia showing distinct behaviours. (C) Box plots showing that none of the parameters measured were significantly different in filopodia expressing mNG-TOCA-1 compared to mNG alone (boxes show median and IQR of mNG-TOCA-1 data, grid lines show median and IQR for control data; see Table S1 for full data). (D) Plotting the log frequency of tip movement velocities reveals a biexponential shape similar to a Laplace distribution. (E) Cross correlation functions (CCFs) for each filopodium at multiple time offsets between tip movement and tip TOCA-1 fluorescence, with hierarchical clustering based on CCFs from −6 to +6 s. Colour shows degree of correlation. Likelihood of the observed level of correlation occuring by chance in individual filopodia indicated by grey/orange boxes to the right, as computed using Markov Chain simulations (see main text). P values shown in the figure are not corrected for multiple comparisons. (F) Averaging the top-correlating, “TOCA-1-responding” sub-cluster (1 in (E), blue) separately from the rest of the filopodia (2 in (E), green) reveals a strong correlation between tip movement and fluorescence, peaking at −2 s time offset. Shading represents 95% confidence interval. A randomisation approach to testing significance (*right*) shows that the observed degree of correlation (blue line) in the TOCA-1-responding subcluster is stronger than would be expected if the two parameters were decoupled (p = 0.014). (G) Measuring fluorescence intensity of mNG-TOCA-1 and GAP43-RFP at the filopodium base (or the predicted base before formation), shows that mNG-TOCA-1 fluorescence peaks around 10 s prior to filopodium emergence and falls away before protrusion begins. Shading represents 95% confidence interval.

To determine whether TOCA-1 had a similar relationship to filopodial protrusions as Ena in our previous observations (Urbančič *et al*., 2017), filopodia from 11 growth cones were analysed in detail using Filopodyan, a semi-automated pipeline for segmenting filopodia and extracting dynamic properties (Urbančič *et al*., 2017). As found previously for Ena, RGCs expressing mNG-TOCA-1 had no significant differences in any of the measured dynamic parameters compared to expression of mNG alone (Fig. 3C, Table S1). In *Drosophila* and filopodia-like structures, filopodia growth and shrinkage rates fall on a bi-exponential Laplace distribution (Dobramysl *et al*., 2021) and we observe a similar relationship here with a slight deviation at the highest velocity protrusion events, which may be because of adhesion to a stiff laminin-coated dish compared to the pliable cells and ECM in native tissues we measured previously (Fig. 3D).

By measuring tip movement speed alongside tip mNG-TOCA-1 fluorescence, a crosscorrelation function (CCF) score for each filopodium at each time point, can be calculated from +1 (perfect positive correlation) to −1 (perfect negative correlation). To ask how frequently mNG-TOCA-1 tip fluorescence correlates with tip extension/retraction, and any offset in peak fluorescence relative to tip movement, CCFs were calculated across a range of time offsets, from −40 to +40 s, followed by hierarchical clustering of the CCFs produced per filopodium (Fig. 3E). A continuum is seen between weak negative correlation, weak positive correlation, and, for a sub-cluster of 16/116 “TOCA-1 responding” filopodia, a strongly significant positive correlation (Fig. 3E,F). The intense redundancy and heterogeneity observed for filopodia is consistent with all our previous observations (Urbančič *et al*., 2017; Dobramysl *et al*., 2021).

For each filopodium 10,000 Markov chain simulations were carried out for tip movement and tip fluorescence separately, and CCFs calculated for each simulation pair, showing that for most of the “TOCA-1 responding” filopodia, the CCFs were significantly higher than expected by chance, based on a Markov chain-based simulation (simulated CCFs exceeded observed in <100/10,000 cases for 9/16; Fig. 3E). The subset of filopodia with the strongest correlations between TOCA-1 intensity and movement are seen with peak correlation at an offset of −2 s prior to forward tip movement (CCF = 0.56, Fig. 3F). To confirm that this was not an artefact of clustering, 1000 randomised datasets were produced by changing the order of the tip movement data (Urbančič *et al*., 2017). CCFs were calculated as before and the TOCA-1 responding cluster was identified in each case, showing that in only 14/1000 cases was the mean CCF higher than the observed dataset (at time offset = 0). The link between mNG-TOCA-1 fluorescence and forward tip movement suggests that TOCA-1 is functionally involved in filopodia extension, similar to Ena and VASP (Urbančič *et al*., 2017).

TOCA-1 forms puncta on the plasma membrane with ^~^4% of these becoming filopodia (Fig. 2A). To track these puncta specifically, we monitored the sites of filopodial initiation on the lamellipodium (Fig. 3G). TOCA-1 fluorescence increases in the −20 to −10 s prior to protrusion and then drops, both before tip emergence (−10 to −2 s) and along with tip protrusion (−2 to 0 s). This is a different pattern to Ena which peaks at −2 s and only drops as the tip emerges (Urbančič *et al*., 2017). This suggests that TOCA-1 plays an early role when filopodia grow from complexes of actin regulatory proteins and is sometimes required for extension of mature filopodia.

### Partial colocalization of TOCA-1 and Ena in filopodia at super-resolution

To understand whether our protein interaction studies *in vitro* were recapitulating a TOCA-1/Ena interaction in filopodia, we expressed mNG-Ena in RGCs alongside immunostaining for TOCA-1. It is clear that while TOCA-1 and Ena puncta are often distinct, many filopodia shafts and tips contain both TOCA-1 and Ena (Fig. 4A). We quantified the areas of overlap finding typically 1-3 per filopodium and, similarly to our results with TOCA-1 immunostaining alone, the density of overlapping puncta were fewer in longer filopodia (Fig. 4B). We speculated that these puncta might be endocytic vesicles (Nozumi *et al*., 2017; Gallop, 2020). Two-colour single molecule localisation microscopy offers an opportunity to examine protein-specific ultrastructure and co-localisation at ^~^50 nm resolution, so we combined expression of mEos-Ena and photoactivated localisation microscopy (PALM) with immunostaining of TOCA-1 and stochastic optical reconstruction microscopy (STORM) using Alexa Fluor-647-anti-rabbit secondary antibodies. Combined PALM/STORM resolves filopodia tips at high resolution, confirming that Ena and TOCA-1 are juxtaposed at a portion of filopodia tips (Fig. 4C, i-iii), while no consistent organisation or architecture, such as vesicular morphology, was evident, potentially due to low protein abundance. TOCA-1 and Ena coincide in places at the leading edge (Fig. 4C, v-vi), possibly corresponding to sites of filopodial formation, though actively extending filopodia are difficult to identify in the absence of a specific marker. Together, these data are consistent with a portion of Ena and TOCA-1 engaging in a transient interaction that particularly occurs in short or newly forming filopodia.

**Fig. 4.**
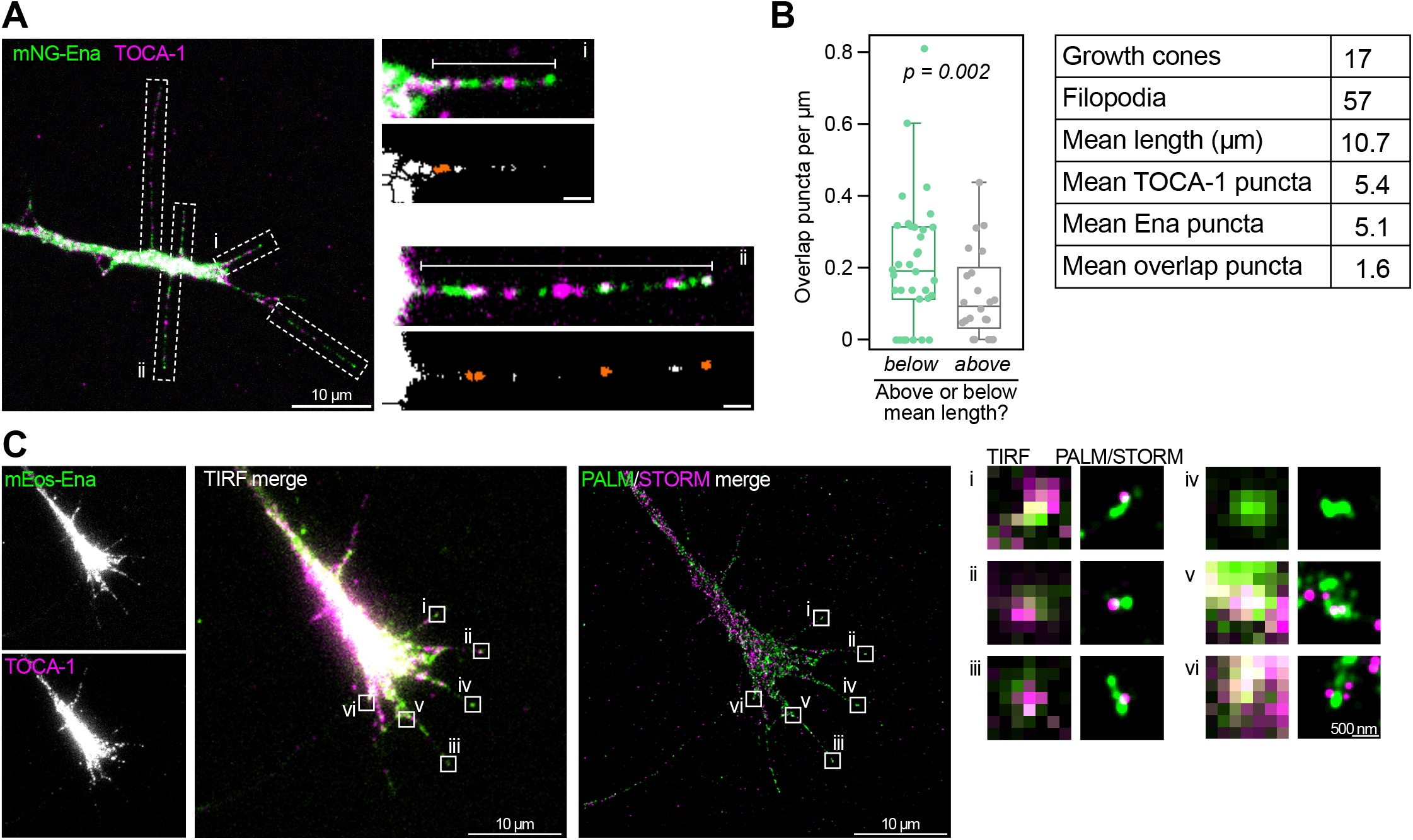
TOCA-1 and Ena do not consistently overlap in fixed cells. (A) Filopodia in fixed RGCs expressing mNG-Ena and immunostained against TOCA-1 show that both form multiple, distinct puncta along filopodia, sometimes at bases (i) and shafts, and sometimes at tips (ii), with systematic manual quantification (white dashed boxes, with examples (i) and (ii) showing areas of TOCA-1/Ena overlap after thresholding, in orange) showing the number of overlapping puncta is around 30% of either Ena or TOCA-1 puncta numbers. (B) Grouping filopodia by size reveals that shorter filopodia have a higher density of TOCA-1 and Ena overlap puncta (p = 0.002; Kruskal-Wallis test). (C) 2-colour PALM/STORM imaging of fixed RGCs expressing mEos-Ena and immunostained against TOCA-1 (TIRF reference images, *left*). This confirms that many filopodia have both Ena and TOCA-1 at tips (features i-iii), others have only one present (iv) while Ena and TOCA-1 also coincide at possible sites of filopodial formation, both lamellipodia (v) and short protrusions (vi). However, there is no consistent arrangement of the two proteins, indicating there is no stable, structured complex. Scale bars 1 μm unless indicated.

### A transient coincidence of TOCA-1 and Ena during filopodial initiation and re-extension

We have demonstrated that increases in TOCA-1 and Ena each precede filopodial initiation and correlate with forward movement in a subset of extending filopodia (Fig. 3) (Urbančič *et al*., 2017) and that TOCA and Ena have a biochemical interaction that is dependent upon a highly clustered state of TOCA-1 (Fig. 1). In cells, such interactions are likely dependent on the Cdc42 activation state, lipid composition of the membrane and competitive interactions with the other interaction partners of both TOCA-1 and Ena (for example N-WASP, Diaph3, IRSp53, lamellipodin). We reasoned that two-colour dynamic imaging would allow us to test whether the partial co-localisation of Ena and TOCA-1 seen in images of fixed cells corresponds to a transient event that is nonetheless predictive of filopodial extension.

We expressed mNG-TOCA-1 with mScarlet-Ena in RGCs and closely examined the rapid dynamics of the protein localisations alongside growth cone morphology and filopodial dynamics (Fig. 5A, Video S4). As well as initiation from the lamellipodium, where Ena and TOCA-1 colocalised, filopodia frequently re-accelerated their protrusion when met by an existing lamellipodium or filopodium. For each event, the tip was tracked manually using TOCA-1, Ena or background fluorescence, and the intensity of mNG-TOCA-1 and mScarlet-Ena was measured at the tip (panel i in Fig. 5B,C,E). The tracking was done manually because use of Filopodyan requires segmentation using a single membrane marker and would require 3 channel simultaneous imaging which is currently beyond our technical capabilities. All filopodia protrusion events (where filopodia tips moved forwards) were classified as either re-extension, if an existing filopodium tip resumed forward movement after stalling (20/37 filopodia from 15 growth cones) or initiation when new filopodia formed directly from the lamellipodium (17/37). Areas of TOCA-1/Ena overlap were identified by setting an intensity threshold for each channel and automatically detecting all areas with both proteins present with a minimum area of 0.06 μm^2^ (approx. 14 pixels, to avoid single pixel fluctuations) and maximum of 1 μm^2^ (panel ii in Fig. 5B,C,E). The presence or absence of overlap at the tip was compared to tip velocity during filopodia protrusion events (panel iii in Fig. 5B,C,E).

**Fig. 5.**
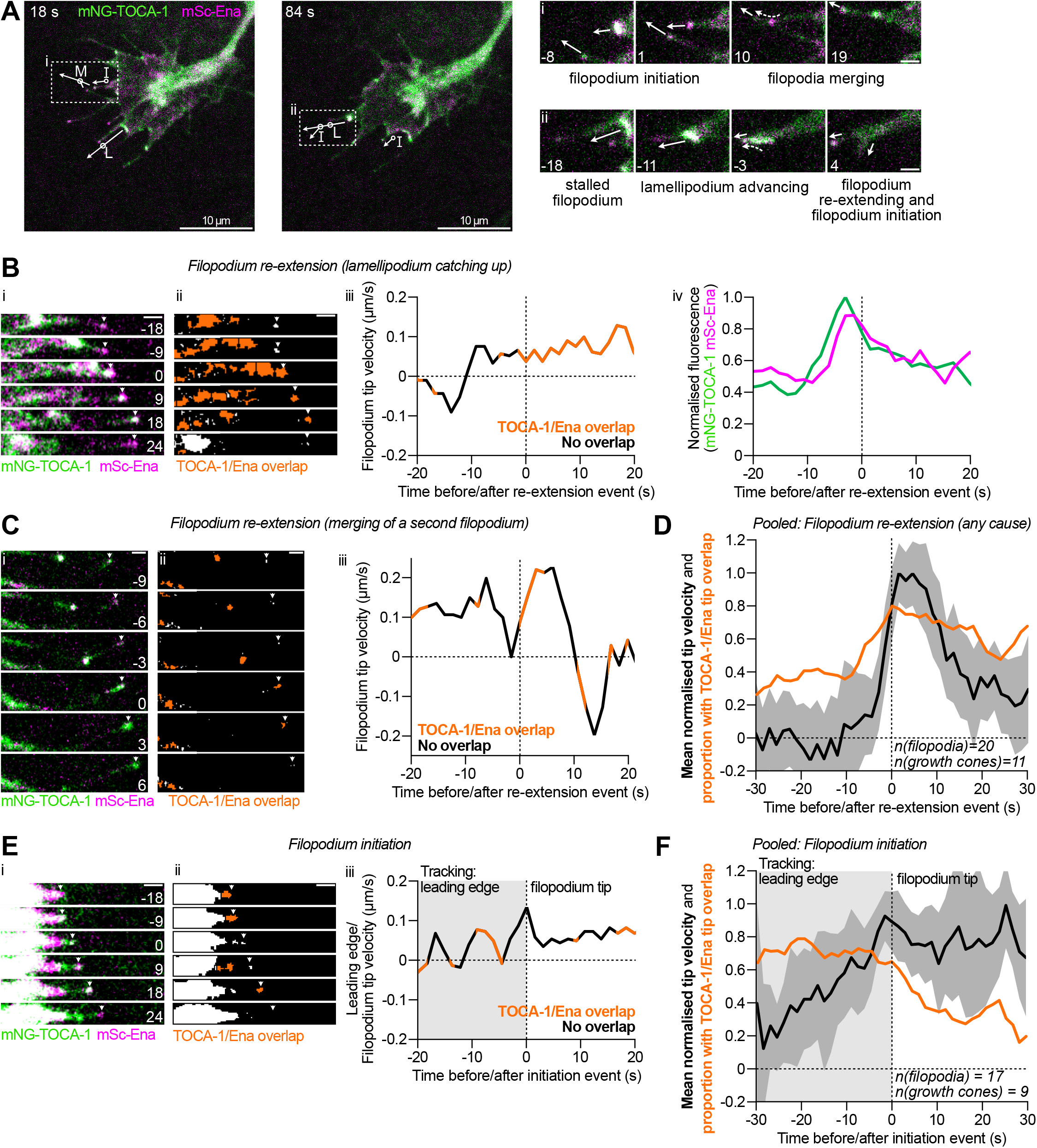
TOCA-1 and Ena coincide transiently during filopodial initiation and re-extension. (A) 37 filopodial initiation/re-extension events were collected from 2-colour, simultaneous TIRF imaging of 15 growth cones (injected with mNG-TOCA-1 and mScarlet-Ena (mSc-Ena); 3 minutes at 1.5 s per frame) and assigned as initiation (17/37) or re-extension (20/37). In these examples (both from Video S4) multiple protrusion events were observed and some filopodia were observed undergoing two types of events. Arrows indicate ongoing/future tip movement for selected protruding filopodia, with circles showing site of protrusion event (I = initiation, L = re-extension after lamellipodium catching up, M = re-extension after merging of a second filopodium). Boxes i and ii are shown expanded (*right*) to show time course of compound protrusion events. Arrows indicate ongoing/future tip movement, dashed arrows indicate movement of fluorescent TOCA-1/Ena puncta towards tips. (B) The tip position was tracked (arrows in B, C, E), and both fluorescence intensity and presence/absence of TOCA-1/Ena overlap were quantified at tips (based on manual thresholding and minimum and maximum areas). In re-extension events caused by the lamellipodium catching up with a static filopodium, (i) the tip was typically high in Ena but low in TOCA-1/Ena overlap before re-extension (−18 and −9 s), then TOCA-1 and Ena levels peaked at or shortly before re-extension (0 and 9 s) and overlap remained high for some time after re-extension (18 s). (ii) Orange puncta in thresholded binary images show areas of TOCA-1/Ena overlap, including large areas of the lamellipodium. (iii) The velocity of the filopodium tip is around zero before re-extension, then accelerates rapidly, coinciding with onset of TOCA-1/Ena overlap (orange line segments). (iv) Measuring the fluorescence intensity of TOCA-1 and Ena at the tip shows a peak shortly before re-extension. (C) In reextension due to merging of a second filopodium, (i) the two tips combine to form a large tip, typically with both TOCA-1 and Ena. (ii) Orange puncta show areas of TOCA-1/Ena overlap. (iii) Velocity of the original filopodium tip undergoes rapid acceleration after merging with the second filopodium, coinciding with transient TOCA-1/Ena overlap. (D) Pooling 20 re-extension events confirms that TOCA-1/Ena overlap levels (orange line) at static tips are low, rising to high levels around re-extension, as seen by the transient burst of high mean tip velocity (black line, shading represents 95% confidence interval). (E) During initiation events, (i) high levels of TOCA-1 and Ena overlap were present at the predicted base on the lamellipodium preceding initiation, and sometimes persisted for a short time after before dropping. (ii) Orange puncta show areas of TOCA-1/Ena overlap. (iii) The velocity of the filopodium tip (black line) peaks around initiation (orange line segments show frames where TOCA-1/Ena overlap at the tip; grey box shows the time period before formation when predicted base was tracked). (F) Pooling 17 initiation events confirms that the prevalence of TOCA-1/Ena overlap at tips (orange line) was high before initiation before gradually falling to low levels, while the mean velocity of the tip peaked around initiation and stayed high (black line, shading represents 95% confidence interval). All scale bars 1 μm unless indicated, time indicated in seconds relative to initiation/re-extension event.

Filopodial re-extension events were most often preceded by an advancing lamellipodium catching up with a static filopodium tip (11/20 events, annotated L, Fig. 5A panel ii and 5B). Frequently, both mNG-TOCA-1 and mScarlet-Ena were present at the lamellipodium, but one or both (especially mNG-TOCA-1) was absent from the static tip (Fig. 5B, panel i). The resumption of filopodium tip protrusion coincided with arrival of the lamellipodium and movement of TOCA-1 and Ena puncta from the advancing lamellipodium to the filopodium tip, with TOCA-1/Ena overlap rare before re-extension and common afterwards (Fig. 5B, panel ii-iii). For both proteins, peak fluorescence typically occurred prior to re-extension, followed by a drop in tip fluorescence especially for TOCA-1 (Fig. 5B, panel iv).

In 6/20 cases, the merging of a second filopodium preceded filopodia re-extension, usually starting with the second tip contacting the first shaft (Fig. 5A panel i, annotated M and 5C, panel i). Again, the overlap of TOCA-1 and Ena puncta was less common before re-extension and more common after, when puncta from the second tip rapidly migrated up the first shaft to join a new enlarged tip (Fig. 5C, panel ii-iii). The original filopodia typically protrude slowly or are stalled before merging, and after merging can show rapid acceleration for some time (10 s in Fig. 5C panel iii).

Pooling all re-extension events reveals that less than 40% of filopodia display TOCA-1/Ena overlap at static filopodia tips from −30 to −8s before re-extension, with filopodia tips often showing a punctum of Ena but not TOCA-1 (Fig. 5D; orange line). The proportion of filopodia showing TOCA-1/Ena overlap rises sharply to greater than 80% of filopodia at the moment of re-extension, gradually falling to 67% around 20 s after. The increase in filopodia with TOCA-1/Ena overlap precedes the increase in mean tip velocity (Fig. 5D; black line), suggesting that a transient TOCA-1/Ena complex is associated with re-starting filopodial extension at mature tips.

During filopodial initiation (Fig. 5A, annotated I, and Fig. 5E, panel i), we tracked the movement and fluorescence at the predicted base (the nearest region on the membrane to where the filopodium will form) before filopodial initiation, then at the tip after initiation. Both TOCA-1 and Ena were abundant at the site of formation before protrusion, and typically both were present at the nascent filopodium tip, but after a few seconds one or both (especially TOCA-1) faded from the tip. To take one example, TOCA-1 and Ena overlap before initiation (−18 and −9 s), and both are present briefly at the nascent tip (18 s, Fig. 5E). This same pattern was observed when pooling multiple examples (Fig. 5F). Overlap of TOCA-1 and Ena at the predicted base was observed in ^~^70% of initiating filopodia at all timepoints during the 30 s before initiation (Fig. 5F). From 2-14 s after initiation, the proportion of filopodia tips with TOCA-1 and Ena overlap steadily falls to ^~^30%, suggesting that the TOCA-1/Ena complex forms only transiently at the plasma membrane before and during the moment of filopodial protrusion. Together, these observations support a model in which TOCA-1 and Ena transiently associate before and during filopodia protrusion events, both initiation and later re-extension.

## Discussion

Untangling the control mechanisms of actin polymerisation at the filopodium tip is difficult because of the level of variability between cell types and cell states and the heterogeneity and redundancy of actin regulators, which may act transiently and play multiple roles. In this study, we show that TOCA-1 localises to filopodia, lamellipodia and inwardly moving puncta in axonal growth cones, and we demonstrate that TOCA-1 and Ena form a new regulatory complex involved in promoting filopodial protrusion, based on transient coincidence before and during filopodium initiation and re-extension.

### TOCA-1 and other membrane adaptors in filopodia formation

TOCA-1 and other F-BAR proteins have well-described roles in endocytosis and promotion of positively curved membrane structures such as cytoplasmic tubular networks (Frost *et al*., 2008; Taylor *et al*., 2019). While the inverse F-BAR proteins SRGAP1-3 bind to negatively curved membrane like I-BAR proteins (Guerrier *et al*., 2009), positively curved F-BAR proteins have also been shown to induce, or localise to, membrane structures with mostly negative curvature such as filopodia (including Gas7 (She *et al*., 2002), PACSIN-1/-2 (Qualmann and Kelly, 2000; Shimada *et al*., 2010), Nwk (Becalska *et al*., 2013; Zhai *et al*., 2022)), suggesting that any membrane curvature preference of F-BAR proteins does not limit them to certain cellular functions. This could be possible because of alternative binding modes that target flat membrane (Frost *et al*., 2008; McDonald *et al*., 2015), or the presence of complex membrane curvatures for example those observed by EM in dendritic filopodial precursors (Galic *et al*., 2014) and described theoretically (Mancinelli *et al*., 2021). Stochastic membrane fluctuations, producing both positive and negative curvature, are proposed to recruit diverse BAR superfamily proteins, with either curvature preference (Mattila *et al*., 2007; Mancinelli *et al*., 2021). Our results showing that TOCA-1 mutants lacking an intact SH3 domain showed slight enrichment to swollen and curved sites (Fig. 2C) is consistent with observations of N-BAR protein ARHGAP44 (Mancinelli *et al*., 2021).

F-BAR proteins in the same sub-family as TOCA-1, CIP4 and FBP17, have multiple opposing roles in mouse cortical neurons, each promoting or opposing neurite formation and endocytosis depending on the isoform expressed (Taylor *et al*., 2019). In these neurons, CIP4 promotes the formation of thin actin ribs that do not extend beyond the leading edge and co-localise with Ena/VASP and formin DAAM1 (Saengsawang *et al*., 2013). As well as F-BAR protein paralogues, other proteins, including lamellipodin (Krause *et al*., 2004), I-BAR proteins like IRSp53 (Disanza *et al*., 2013), and recently formin FMNL2 (Fox *et al*., 2022), have been shown to play comparable roles to that proposed for TOCA-1 as a membrane adaptor and scaffold for Ena/VASP actin filament elongators or other actin regulators. This variety may reflect the different filopodia being studied or alternative mechanistic pathways, fitting our observations that filopodia tips are heterogeneous, with stochastic assembly of one or more redundant sub-complexes of actin regulators being sufficient to drive filopodial formation (Dobramysl *et al*., 2021). It may be that certain actin regulatory activities are required, such as a membrane adaptor scaffold and an actin filament elongator, but that these can be achieved via multiple regulatory pathways (though Myosin-X has been recently reported as uniquely critical for microspike tip VASP clusters in B16-F1 melanoma cells (Pokrant *et al*., 2022)).

IRSp53 appears to localise stably to filopodia shafts (Sudhaharan *et al*., 2019; Cheng and Mullins, 2020; Tsai *et al*., 2022). In contrast, TOCA-1 localises to discrete puncta, especially at filopodial tips (Fig. 2D). The multiple shaft puncta observed by immunostaining of endogenous TOCA-1 may correspond to the dynamic puncta moving up and down shafts observed in videos of mNG-TOCA-1, or a pool of TOCA-1 not labelled by mNG-TOCA-1. Live imaging shows that puncta of TOCA-1 are mostly present at tips or transiently during initiation/re-extension events (Fig. 2A), suggesting a specific role in an actin regulatory complex, perhaps differing from a more structural role for I-BAR proteins in stabilising curved filopodial membrane.

Similar to clusters of lamellipodin, TOCA-1 was first recruited to plasma membrane, then moved laterally and coalesced into larger puncta that recruited other components and sometimes developed into filopodia (Fig. 2B) (Cheng and Mullins, 2020). Furthermore, the frequent observation of TOCA-1 puncta moving inwardly, often associated with filopodial formation, is consistent with size-dependent splitting of the TOCA-1 cluster to maintain appropriate stoichiometry, as observed for the lamellipodin-VASP complex (Cheng and Mullins, 2020), though other explanations are possible such as coincident retrograde movement of a membrane vesicle (Nozumi *et al*., 2017; Gallop, 2020)

### The mechanisms of TOCA-1 promoting filopodia protrusion

Though TOCA-1 and Ena interact directly *in vitro* (Fig. 1D), they did not persistently co-localise in these cells, suggesting the conditions needed for interaction are rare, be it sufficient density of TOCA-1 or the influence of other binding partners. Fixed imaging did not capture a consistent arrangement of TOCA-1 and Ena, even at super-resolution (Fig. 4C), though rapid, time-resolved super-resolution imaging could be useful in revealing any arrangement of the transient TOCA-1/Ena complex during filopodial protrusion and how it links to the membrane.

We showed that TOCA-1 correlates with filopodia protrusion (Fig. 3E), and that TOCA-1 and Ena overlap peaks transiently around re-extension events (Fig. 5D), confirming that it is functionally relevant at mature tips. The proportion of filopodia tips with TOCA-1 and Ena overlap was substantially higher before/during filopodium initiation and re-extension compared to mature tips (after initiation or before re-extension). Measuring the degree of overlap is a simplification to capture and quantify the coincidence of the two proteins, but where fluorescence intensity of each protein was quantified (Fig. 5D, part iv), it fits with a transient peak in abundance associated with dynamic filopodia events. Use of Split-GFP may prove helpful in future studies to monitor the direct interaction in cells.

As well as filopodial extension, TOCA-1 appears to have a particular role in filopodial initiation, since it arrives early (before Ena) at sites of filopodial initiation (Fig. 3G), and both TOCA-1 itself (Fig. 2E) and TOCA-1/Ena overlap (Fig. 4B) are more abundant at the tips of short (potentially young) filopodia. N-WASP, an activator of Arp2/3 complex and branched actin, binds TOCA-1 at high and low densities (Fig. 1F). It is possible that when TOCA-1 is initially recruited to leading edge membranes, at lower density, it promotes polymerisation of branched actin structures via N-WASP or WAVE, such as advancing lamellipodia. Then, after attaining sufficiently high density by coalescence of multiple puncta, TOCA-1 may switch to promoting linear actin polymerisation via Ena, leading to filopodial initiation or re-extension. The clustering of TOCA-1 required for Ena binding is similar to observations with VASP clustered on beads (Breitsprecher *et al*., 2008) and the increased processivity similar to the activity of Ena on filaments clustered by fascin (Harker *et al*., 2019).

Further work is needed to explore other candidate downstream actin regulators involved in TOCA-1-mediated filopodial protrusion, such as Myosin-X which has been implicated in multi-cycle filopodial extension (He *et al*., 2017), and the possible interplay between the TOCA-1/N-WASP/WAVE and TOCA-1/Ena complexes, confirming if they are spatially and temporally separated, and dissecting the relationship between advancing lamellipodia and protruding filopodia.

## Figure Legends

**Fig. S1. TOCA-1 localises to lamellipodia and filopodia in human neuroblastoma SH-SY5Y cells.**

(A, B) Montages showing transfected human mNG-TOCA-1 (green) forming puncta at advancing lamellipodia (arrows) and filopodia tips (arrowheads). F-actin labelled with transfected F-tractin-tdTomato probe. Time indicated in seconds.

**Fig. S2. Affinity purified rabbit anti-TOCA-1 antibody recognises TOCA-1 in lysate and cells.**

(A) SDS-PAGE gel with Xenopus egg extract (HSS) or purified SNAP-TOCA-1 (TOCA), stained with the indicated bleeds at 1:500 dilution, showing that before immunisation, a small nonspecific band was present with concentrated, purified SNAP-TOCA-1 and not with HSS. All post-immunisation bleeds recognise TOCA-1 in HSS and purified SNAP-TOCA-1, and after affinity purification (“purified harvest”) the specificity is greatly improved. (B) RGCs expressing membrane marker GAP43-RFP. No primary antibody immunostaining control, showing that fluorescence is specific to anti-TOCA-1/anti-rabbit-488. Contrast applied equally between images for 488 channel and allowed to vary in GAP43-RFP channel to account for variable expression levels. (C) RGCs expressing mNG-TOCA-1, immunostained with anti-TOCA-1 / anti-rabbit-AF647 and phalloidin-AF568, showing a similar pattern of fluorescence between endogenous and exogenous TOCA-1. (D) Acquiring images with no, one or both lasers active confirmed that almost no fluorescent signal in the 568 channel was due to excitation of either fluorophore with the 491 nm laser, and vice versa. No background subtraction, contrast applied equally to all eight images.

**Table S1. Filopodia morphological parameters from Filopodyan, showing no difference between mNG alone and mNG-TOCA-1.**

**Video S1. RGC growth cone expressing mNG-TOCA-1 and GAP43-RFP, showing TOCA-1 recruitment to filopodia, lamellipodia and inwardly-moving puncta.**

**Video S2. RGC growth cone expressing mNG-TOCA-1 (green) and GAP43-RFP (magenta), showing detail of filopodium formation, with two puncta of mNG-TOCA-1 moving laterally on the plasma membrane and coalescing before initiation. Image de-noised with nd-safir. Time relative to filopodium formation, scale bar 1 μm.**

**Video S3. RGC growth cone expressing mNG-TOCA-1-SH3^W517K^ and GAP43-RFP, showing no enrichment of TOCA-1 to filopodia, but transient enrichment to sites of swollen and curved membrane.**

**Video S4. RGC growth cone filopodium expressing mNG-TOCA-1 and mScarlet-Ena, showing that the two proteins coincide during filopodial protrusion events (I = initiation, M = re-extension after merging, L = re-extension after lamellipodium catching up with filopodium).**

## Materials and Methods

### Plasmids

pET-His-SNAP-TOCA-1 (*Xenopus tropicalis*; BC080964), pCS2-His-SNAP-Ena (*X. laevis*; BC073107) and pET-KCK-VASP (*X. laevis*; BC077932), or SNAP alone, were generated previously (Dobramysl *et al*., 2021). mNeonGreen was supplied by Allele Biotechnology and Pharmaceuticals (Shaner *et al*., 2013) and pCS2-mNG-Ena and pCS2-mNG alone were generated previously (Urbančič *et al*., 2017). New vectors were generated by PCR (Phusion-HF, NEB) into parent vectors digested with FseI/AscI unless otherwise stated. pCS2-mNG-TOCA-1 was generated by sub-cloning TOCA-1 into the digested pCS2-mNG vector, with oligonucleotide primers: 5’-GCATGGCCGGCCACCATGAGCTGGGGTACTG-3’ and 5’-GGCGCGCCTTAGATATAAGTTACTGC-3’. Ena was subcloned using primers: 5’-GCATGGCCGGCCACCATGAGTGAACAGAGCATC-3’ and 5’-GGCGCGCCCTATGCGCTGTTTG-3’ into pCS2 vectors generated with mEos3.2 (amplified from pmEos3.2-N1, which was a gift from Michael Davidson & Tao Xu (Zhang *et al*., 2012); Addgene plasmid 54525) or mScarlet (amplified from pLifeAct_mScarlet_N1, which was a gift from Dorus Gadella (Bindels *et al*., 2017); Addgene 85054). GAP43-RFP was a gift from the Holt laboratory. F-tractin was a gift from J. A. Hammer III. A version of His-Ena without SNAP was generated using the above Ena primers. Ena-ΔPRR was generated by replacement of S291-G373 with a linker sequence GGGGSSGG, using In-Fusion cloning (Takara Bio) with primer pairs: 5’-TCATCATCACGAATTCAGGCCGGCC-3’ + 5’-ACCTGAAGAACCACCTCCTCCCACTCTCCGTTCCCTTTCCCATTCC-3’ and 5’-GGTGGTTCTTCAGGTGGATCAGAAGAGAATCGTGCTTTATC-3’ + 5’-GGCCGCGGCGCCAATGCATTGGGCC-3’ into the parent vector digested with EcoRI and NotI. VASP-ΔPRR was generated by removal of S116-S192, using In-Fusion cloning with primer pairs: 5’-ACAATTCCCCTCTAGAAATAATTTTG-3’ + 5’-CACCCCCACCAGTCTCCAGTGCATCCAAGG-3’ and 5’-AGACTGGTGGGGGTGGAGGAAGCTCAGGTGG-3’ + 5’-TATCATCGATAAGCTTTAATGCGGTAG −3’ into the parent vector digested with HindIII and XbaI. The additional mutation in construct M1 (P234G) was generated by PCR (Pwo Master; Roche) using primers: 5’-CCTCCCCAGTTGGTGGAGTGGGTGCAAAGCCAGACATAAGTCG-3’ and 5’-CGACTTATGTCTGGCTTTGCACCCACTCCACCAACTGGGGAGG-3’. TOCA-1 mutants were generated as shown in Fig. 1A. For mutants starting at the N-terminus, the forward primer was 5’-GCATGGCCGGCCACCATGAGCTGGGGTACTG-3’ and the reverse primers were 5’-GGCGCGCCTTAGCTGTAGTCTTCAAAGGGATAGTC-3’ (FBAR only), 5’-GGCGCGCCTTATTGTGCTACAAGATGGTTAGCTTC-3’ (FBAR-HR1) and 5’-GGCGCGCCTTAAGCTGGGAGAGGTTCATCATC-3’ (FBAR-HR-linker). The SH3 only mutant was generated using primers: 5’-GCATGGCCGGCCACCATGGGACACTGCAAATCAC-3’ and 5’-GGCGCGCCTTATAGAGTGATATCTATGTAGGATGTGG-3’ and the W517K mutant was generated using primers: 5’-GATAAAGGGGATGGAAAGACAAGAGCAAG-3’ and 5’-CTTGCTCTTGTCTTTCCATCCCCTTTATC-3’. Human TOCA-1 was a gift of Kwonmoo Lee and was sub-cloned into the pCS2-mNG vector using primers 5’-GATCGGCCGGCCATGAGCTGGGGCACGGAGC-3’ and 5’-GGCGCGCCCTGCAGCTCGAGTCAGGAACC-3’.

### Protein expression and purification

His-tagged proteins were expressed and purified with Ni-NTA columns and gel filtration on S200 columns as previously (Dobramysl *et al*., 2021) with some exceptions. His-SNAP-TOCA-1 was purified in high salt buffers (300 mM NaCl instead of 150 mM NaCl during washing and elution), with two additional washes with 50 mM imidazole elution buffer before elution in a single 300 mM imidazole step and concentration using a spin concentrator before proceeding to gel filtration. TOCA-1 mutants were expressed and purified in the same way as wild type TOCA-1. His-Ena-ΔPRR was expressed in 293F cells and purified in the same way as wild type Ena (Dobramysl *et al*., 2021), except that 300 mM NaCl, not 150 mM, was used in wash and elution buffers. His-KCK-VASP mutants were expressed in BL21 *E. coli* in the same way as Ena mutants. His-SNAP alone was purified with elutions at 100 mM and 300 mM imidazole.

### Coupling to beads, including densities, and precipitation from HSS

For each reaction, 20-40 μl SNAP-Capture beads (NEB, S9145S) were pre-equilibrated in 150 mM NaCl, 20 mM HEPES (pH 7.4), 0.1% TWEEN-20, then 500 μl of SNAP-coupled protein was added overnight, under rotation at 4°C in buffer containing 150 mM NaCl, 20 mM HEPES (pH 7.4), 0.1% TWEEN-20 and 1 mM DTT. Beads were washed five times in 150 mM NaCl, 50 mM Tris, 1 mM DTT, 0.1% TWEEN-20. The capacity of benzylguanine sites on beads was determined empirically for each protein, with, per 40 μl of beads, 500 μl of 12 μM SNAP-TOCA-1 or mutants; 500 μl of 24 μM SNAP alone. For varying the density of TOCA-1 on beads, 500 μl of 2 μM SNAP-TOCA-1 was coupled to 10 μl of beads (determined to be maximum capacity) or 50 μl (intermediate density) or 100 μl (low density), then, respectively, 90 μl, 50 μl or 0 μl of un-coupled beads were added to make a final volume of 100 μl beads for each condition, and a control sample with 100 μl un-coupled beads was also prepared. In a second stage of coupling, 500 μl of 80 μM SNAP alone was then added, to bind the remaining benzylguanine sites on the beads. To precipitate proteins from *Xenopus* HSS, coupled beads were incubated for 1 h at 4°C with 400 μl *Xenopus* HSS (prepared as previously described (Walrant *et al*., 2015) and diluted to 4.17 mg/ml in 50 mM Tris, 150 mM NaCl, 2 mM DTT and energy mix [50 mM phosphocreatine, 20 mM Mg-ATP (adjusted to pH 7.0 with Tris base), 20 mM MgCl_2_]). To precipitate purified Ena/VASP, 100 μl of 1 μM protein was incubated with 10 μl of coupled beads.

### Antibody affinity purification

Affi-Gel 15 beads (Bio-Rad, 1536051) were equilibrated in 300 mM NaCl, 20 mM Na-HEPES pH 7.4, 2 mM EDTA and 2 mM DTT, then incubated with His-SNAP-TOCA-1, or SNAP alone, for 4 h, 4°C under rotation. 1 M mono-ethanolamine pH 8 was added to block (1 h), then the TOCA-1-coupled beads were washed on a column with the following buffers: 500 mM NaCl and 20 mM Na-HEPES, glycine-HCl pH 2.5, tri-ethylamine pH 11.5. The harvest bleed serum was passed through the column coupled to SNAP alone, then the flow through applied to the column coupled to His-SNAP-TOCA-1 (2 h, room temperature, under rotation). The column was washed in 20 ml of 400 mM NaCl, 30 mM Na-HEPES pH 7.7 and 3 ml of 300 mM NaCl, 10 mM Tris-HCl pH 7.2., before elution. Elution with acid was carried out with 100 mM glycine pH 2.5, 300 mM NaCl and neutralised in 1.5 M Tris-HCl pH 8.8, followed by elution with base using 100 mM triethylamine pH 11.5, 300 mM NaCl and neutralisation in 2 M Tris-HCl pH 6.5. Elution fractions were screened by absorbance at 280 nm and pooled, before exchanging buffer to 10 mM K-HEPES, 100 mM KCl, 1 mM MgCl_2_, 100 nM CaCl_2_, pH 7.4 overnight.

### Western blotting and antibodies

Samples for western blotting were separated on 4-20% gradient polyacrylamide gels (Mini-PROTEAN TGX, Bio-Rad, 456-1096) and transferred to nitrocellulose membranes by wet transfer in 25 mM Tris, 192 mM glycine, 0.1% SDS, 20% methanol for 1 h at 0.38 A (Bio-Rad Mini Trans-Blot Cell apparatus) or dry transfer (programme 0, iBlot 2, ThermoFisher). Membranes were blocked in 5% milk powder/0.1% TWEEN-20/Tris-buffered saline (20-60 minutes, room temperature) and stained with primary antibody in blocking solution (1 h at room temperature or 4°C overnight). Membranes were washed 3-5 times in 0.5% milk powder/0.1% TWEEN-20/Tris-buffered saline for 5-10 minutes then incubated with goat anti-rabbit 800CW secondary antibody (Li-Cor, 926-32211; 30-60 minutes, room temperature) before washing as before and imaging on a LI-COR BioSciences Odyssey Sa scanner.

Antibodies for blotting: affinity purified anti-TOCA-1 antibody (described above) or other unpurified bleeds (1:500 dilution). The anti-Ena (1:15000), anti-VASP (1:500) and anti-N-WASP (1:2000) antibodies were affinity purified and described previously (Dobramysl *et al*., 2021), as was the anti-Diaph3 antibody (1:1300), a gift from Marc Kirschner (Ho *et al*., 2004). Antibodies for immunostaining cells: affinity purified anti-TOCA-1 antibody (described above; 1:500 dilution). Secondary antibodies used for immunostaining: goat anti-rabbit-AlexaFluor 488 (1:2000, Invitrogen, A11008), goat anti-rabbit-AlexaFluor 647 (1:2000, Invitrogen, A21244).

### RGC preparation and injection of RNA

This research was regulated under the Animals (Scientific Procedures) Act 1986 Amendment Regulations 2012 following ethical review by the University of Cambridge Animal Welfare and Ethical Review Body. *Xenopus* embryos were fertilised *in vitro*, RNA was introduced by electroporation at stages 26-28 and RGC explants were taken at stages 35-36 and cultured for 19-24 h in 60% L-15 (Sigma-Aldrich, L1518) in water, on 35 mm glass-bottom dishes (MatTek P35G-1.5-14-C) coated with 10 μg/ml poly-L-lysine for 1 h (Sigma, P8920) and 10 μg/ml laminin for 5-10 minutes (Sigma, L2020) as described previously (Falk *et al*., 2007; Leung and Holt, 2008; Urbančič *et al*., 2017). For experiments with TOCA-1 mutants, mScarlet-Ena, mEos-Ena or for immunostaining, 75 pg of RNA was micro-injected of into the neural fated blastomeres of 4 cell embryo instead of electroporation. mNG-TOCA-1 and mScarlet-Ena were co-injected at a ratio of 1:2 to equalise the resultant fluorescence levels. Capped RNA was synthesised after linearisation with NotI using SP6 mMessage mMachine kit (Invitrogen, AM1340) with elution into RNase-free water.

### Live imaging of RGCs

Live imaging of RGCs was conducted in 60% L-15/water under HILO illumination on a custom made TIRF setup described previously (Urbančič *et al*., 2017) with an iLas2 illuminator (Roper Scientific), an Optosplit beam splitter (Cairn Research) and a CMOS camera (Hamamatsu ORCA-Flash4.0). Images were acquired at a rate of 2 s per time point (mNG-TOCA-1/GAP43-RFP videos) or else 1.5 s per time point, with a 100x 1.49 NA oil immersion objective (pixel size 0.065 μm) at room temperature, controlled by MetaMorph software (Molecular Devices).

### Fixed imaging of RGCs and PALM/STORM

RGCs were washed once in 60% L-15/water then fixed in 4% PFA/7.5% sucrose/PBS (0.5 − 1 h, room temperature) and washed three times in 0.002% Triton X-100/PBS. Cells were permeabilised in 0.1% Triton X-100/PBS (3 minutes, room temperature) then washed twice as before and blocked in 5% goat serum/0.002% Triton X-100/PBS (overnight, 4°C). Primary antibody was diluted in blocking solution and added to cells (1 h, room temperature) with three washes in 0.5% goat serum/0.002% Triton X-100/PBS (5 minutes each). Secondary antibody was diluted in blocking solution and added to cells (30 minutes, room temperature) with phalloidin-AlexaFluor568 (1:100 dilution, Invitrogen, A12380) included when indicated, before washing as before. Cells were imaged in wash buffer by TIRF (as for live imaging), or for images in Fig. 2D and Fig. S2C, imaged with a Photometrics Evolve Delta EM-CCD camera instead.

For PALM/STORM imaging, wash buffer was replaced with 150 μl STORM buffer (enzyme mix (50 μg/ml catalase (Sigma), 50 mM Tris-HCl (pH 7.5), 0.5 mg/ml glucose oxidase (Sigma)), 100 mg/ml D-Glucose (Sigma)/ddH_2_O, 100 mM cysteamine hydrochloride (MEA, Sigma, M6500)/ddH_2_O), and dishes were sealed by lowering a coverslip (18 × 18 mm) onto the central well. Images were acquired on an N-STORM system controlled by NIS Element AR version 4.50, with an Agilent laser bed (405 nm, 488 nm, 561 nm, 647 nm lasers), CPI Plan Apo 100x 1.49 NA objective and an N-STORM QUAD filter (405/488/561/647), and imaging with an iXon Ultra 897 EM-CCD camera (Andor). PALM/STORM images were acquired sequentially, in TIRF, with around 10,000 frames (20 ms) of PALM imaging (using low laser power (1-10%) 405 nm to sparsely photoconvert mEos and 561 nm laser at high laser power for imaging) followed by 20,000-30,000 frames (20 ms) of STORM imaging (using low laser power 405 nm to tune blinking rates and 1-2 kW/cm^2^ 647 nm illumination). TIRF reference images (Fig. 4C) were acquired using the same system.

### Culture and imaging of SH-SY5Y neuroblastoma cells

SH-SY5Y cells were a gift from Rick Livesey and were cultured in DMEM (high glucose, Gibco, D6546) with 2 mM GlutaMAX (Gibco, 35050038), 10% FBS, 100 U/ml penicillin and 100 μg/ml streptomycin. 24,000 cells were plated onto the central wells of 35 mm glass bottom dishes coated with laminin (10 μg/ml), then 24 h after seeding media was changed to differentiation media (the same media with 1% FBS and 10 μM retinoic acid) to induce neuronal differentiation and fresh differentiation media was added every 24 h. On day 4 after seeding, cells were transfected with mNG-humanTOCA-1 and F-tractin-tdTomato actin probe using JetPRIME (Polyplus), and cultured with further daily media changes for 48 h, when media was changed to L-15 live imaging media. After 4 h at 37°C, cells were imaged in TIRF as above.

### Image processing and analysis

Image processing was performed in FIJI (Schindelin *et al*., 2012) with custom macros for analysis and some processing macros developed by Steve Rothery at the FILM facility, Imperial College London (www.imperial.ac.uk/medicine/facility-for-imaging-by-light-microscopy/software/fiji/). Images were processed by overlaying the two channels (when Optosplit used) and de-noising with nd-safir (for Fig. 2B, Video S2) (Boulanger *et al*., 2010) or a 50-70 px rolling ball background subtraction for all other images.

For immunostained RGCs, individual filopodia of length ≥3 μm were extracted using the rotated rectangle tool, then measured using a straight line from tip (defined by GAP43-RFP where present, else by the furthest punctum of Ena or TOCA-1 in line with the shaft) to base (defined as the start of a region of consistent, narrow width). Puncta were counted automatically, after manual thresholding to exclude most of the noise, as particles between 0.06 – 1.00 μm^2^ (corresponding to around 14 pixels, or a box of sides 250 nm). Overlap puncta were identified using a binary addition of the two thresholded images, and again automatically selecting resultant overlap puncta with area 0.06 – 1.00 μm^2^. Tip localised puncta were defined as being at least half within a 1 μm circle anchored to the tip. Significance was assessed with a Student’s t-test, if normally distribution according to a Jarque-Bera test, or a Kruskal-Wallis test if not.

Videos of mNG-TOCA-1 and GAP43-RFP-expressing RGCs were analysed with Filopodyan (Urbančič *et al*., 2017) (without de-noising), using thresholding parameters: RenyiEntropy / Fit tip / ED iterations: 4 / LoG sigma 2.6-3.6 depending on the brightness of the video; and filter settings: Min start frame: 1 / Min frames: 3 / Min max length: 1.8 / Min length change: 0.1 / Max mean waviness: 0.38. Tracks assigned to filopodia that merged, moved out of focus or were otherwise poorly annotated were manually excluded. Data tables were then analysed using FilopodyanR, with tip fluorescence processed by normalisation to growth cone body fluorescence and, for Fig. 3E, background subtraction based on signal near the growth cone boundary (Urbančič *et al*., 2017).

TOCA-1 puncta behaviour was quantified from mNG-TOCA-1/GAP43-RFP videos after de-noising. Scoring of numbers of coalescing puncta used distinct puncta that could be tracked throughout the < 6 s before filopodial initiation. For quantifying the fate of TOCA-1 puncta, a macro was used to track all puncta with minimum 5 frames, minimum 0.65 μm link distance and starting within 1 μm of the leading edge. Then individual puncta were randomly selected from a list to then be manually assigned to a category.

Videos of mNG-TOCA-1 and mSc-Ena were processed manually, with background subtraction, extraction of filopodia that underwent initiation or re-extension events using a rotated rectangle to re-orientate parallel to the long axis of the filopodium. Fluorescence intensity values were extracted along the length of the filopodium (averaging across a 20 px column for each pixel along the long axis) using a line profile tool (line/time macro; FILM facility). Overlap puncta were defined as areas of overlap 0.06 – 1.00 μm^2^ after thresholding and binary addition as above, and scored as positive if any pixels in the 20 pixel column were part of an overlap punctum. The tip was tracked manually using TrackMate (Tinevez *et al*., 2017), and a custom Excel worksheet was used to extract the fluorescence intensity values and presence or absence of any TOCA-1/Ena overlap at the tip (averaged over 5 pixel window centred on tip along the long axis of filopodium). The velocity was calculated as displacement along the long axis for each frame and velocity and fluorescence intensity values, and proportion with TOCA-1/Ena overlap, were smoothed by a moving average over 3 frames.

### Super Resolution Image Reconstruction

Image data stacks were converted from Nikon image files into TIF stacks using Fiji. Single molecule blinking events were detected in unprocessed camera frames and fit with a two-dimensional (2D) gaussian model as previously described (Li *et al*., 2018). Fit results were filtered based on number of photons (50 – 5000), localization precision (0.5 – 50 nm), goodness of fit (LLR < 150) and PSF width (sigma, 50-150 nm). Post processing drift correction was applied using a redundant cross-collection algorithm as previously described (Wang *et al*., 2014). Because PALM and STORM datasets were collected sequentially, in that order, drift correction was applied relative to the last frame of the PALM image and to the first frame of the STORM image. Images were reconstructed with a 16 nm pixel size and blurred with a 2D Gaussian equal to the image’s average localization precision.

## Acknowledgements

We would like to thank Asha Dwivedy for help with eye primordia electroporation and dissection, Ken Li for help with analysing the distribution of filopodial growth and shrinkage rates and Jonathan Gadsby for helping with affinity purification of the anti-TOCA-1 antibody.

## Competing Interests

The authors declare no competing interests.

## Funding

This work was supported by Wellcome Trust Research Career Development Fellowship WT095829MA and Senior Research Fellowship 219482/Z/19/Z to JLG and studentship 099740/Z/12/Z to HMF. AW was supported by a Biochemical Society Summer Vacation Studentship. We acknowledge core funding by the Wellcome Trust (092096) and Cancer Research UK (C6946/A14492).

## Data availability

Data are available on request to JLG.

## Notes

### Competing Interest Statement

The authors have declared no competing interest.

