## Supplementary material for "Filopodial protrusion driven by density-dependent Ena-TOCA-1 interactions": Fig. S1

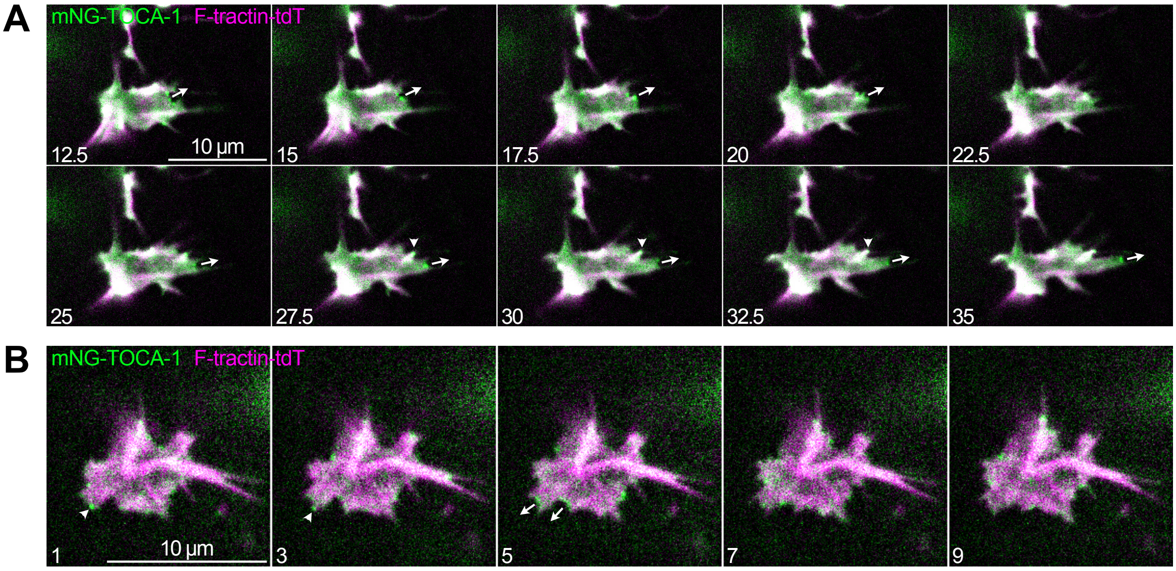


**Fig. S1. TOCA-1 localises to lamellipodia and filopodia in human neuroblastoma SH-SY5Y cells.**
(A, B) Montages showing transfected human mNG-TOCA-1 (green) forming puncta at advancing lamellipodia (arrows) and filopodia tips (arrowheads). F-actin labelled with transfected F-tractin-tdTomato probe. Time indicated in seconds.
