## Supplementary material for "Filopodial protrusion driven by density-dependent Ena-TOCA-1 interactions": Fig. S2

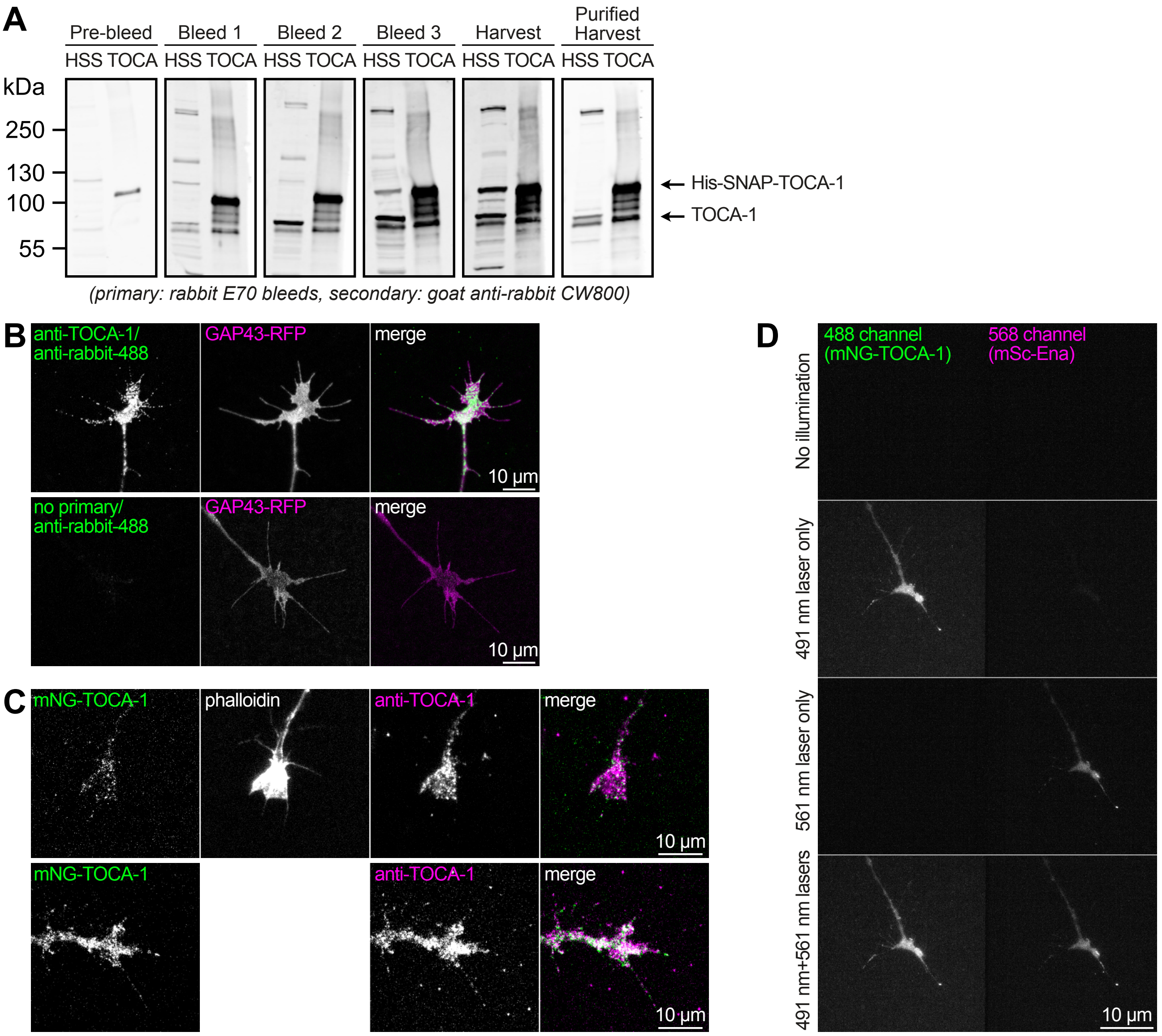


**Fig. S2. Affinity purified rabbit anti-TOCA-1 antibody recognises TOCA-1 in lysate and cells.**

(A) SDS-PAGE gel with *Xenopus* egg extract (HSS) or purified SNAP-TOCA-1 (TOCA), stained with the indicated bleeds at 1:500 dilution, showing that before immunisation, a small non-specific band was present with concentrated, purified SNAP-TOCA-1 and not with HSS. All post-immunisation bleeds recognise TOCA-1 in HSS and purified SNAP-TOCA-1, and after affinity purification ("purified harvest") the specificity is greatly improved. (B) RGCs expressing membrane marker GAP43-RFP. No primary antibody immunostaining control, showing that fluorescence is specific to anti-TOCA-1/anti-rabbit-488. Contrast applied equally between images for 488 channel and allowed to vary in GAP43-RFP channel to account for variable expression levels. (C) RGCs expressing mNG-TOCA-1, immunostained with anti-TOCA-1 / anti-rabbit-AF647 and phalloidin-AF568, showing a similar pattern of fluorescence between endogenous and exogenous TOCA-1. (D) Acquiring images with no, one or both lasers active confirmed that almost no fluorescent signal in the 568 channel was due to excitation of either fluorophore with the 491 nm laser, and vice versa. No background subtraction, contrast applied equally to all eight images.
